# Superresolution imaging of live samples by centroid reassignment microscopy

**DOI:** 10.1101/2025.08.03.668310

**Authors:** Chuan Li, Quan Le, Julian O. Kimura, Bingying Zhao, Yunzhe Li, Jian Zhao, Thomas Bifano, Brandon Weissbourd, John T. Ngo, Jerome Mertz

## Abstract

Superresolution imaging has become one of the most important recent advances in microscopy development. However, most superresolution methods are ill-adapted for live-sample imaging because they are unacceptably slow, susceptible to artifacts, or require the use of specialized fluorophores and labeling protocols. We introduce a superresolution method called centroid reassignment microscopy (CRM) that overcomes these limitations. CRM is a simple variation on confocal microscopy wherein the single-element detector and small pinhole are replaced by a centroid detector and larger pinhole. Superresolution is obtained by reassigning the detected fluorescence to its centroid location of as a function of the scanning excitation focus location. Our method bears resemblance to the method of image scanning microscopy, which involves the use of an array detector, with the advantage that CRM provides improved resolution for the same number of detected photons while being simpler to implement and less sensitive to noise. CRM is light-efficient, fast (single-frame), robust to defocus aberrations, and requires no changes whatsoever in standard fluorescence imaging protocols, making it uniquely attractive for superresolution imaging of live, dynamic samples.

## INTRODUCTION

High-resolution optical imaging is essential for uncovering the dynamic architecture of living biological systems, including mitochondrial fission and fusion^1,2^, axonal transport^3,4^, synaptic plasticity^5,6^, cytoskeletal remodeling^7,8^, etc.. Capturing such events requires imaging approaches that not only deliver high spatial resolution beyond the diffraction limit (< 200 nm) but also maintain long-term sample viability^9^. While several superresolution approaches have been applied to live-sample imaging^9^, these have been hindered by significant challenges. Single-molecule localization microscopy (SMLM) approaches^10–18^ require the use of specialized blinking or photoswitchable molecules, often requiring hundreds or thousands of raw images and restrictive sample preparation protocols. Stimulated-emission depletion approaches^19^ require complex instrumentation involving high illumination powers, with a limited range of fluorophore compatibility. While structured illumination approaches overcome many of these issues, they too suffer from their own drawbacks. Structured illumination microscopy^20,21^ based on frequency multiplexing, requires an accurate estimation of the illumination patterns, making it susceptible to noise and aberrations. Random illumination microscopy^22^, which requires hundreds of raw image acquisitions, is susceptible to motion artifacts. To date, superresolution microscopy approaches have come nowhere close to the broad range of applicably for live-sample imaging as traditional confocal microscopy.

Indeed, traditional confocal microscopy can itself be considered a structured illumination approach^23^, though with limited superresolution capacity. To obtain the best resolution from a confocal microscope, its pinhole must be closed to zero size, which is not possible in practice, thus imposing a compromise between resolution and signal strength. This compromise has been resolved by image scanning microscopy^23,24^ (ISM), which provides essentially the same resolution as a closed-pinhole confocal, but with the light collection efficiency of an open pinhole. ISM involves replacing the single-element detector of a confocal microscope with an array detector, allowing the possibility of superresolution by pixel reassignment. Initial incarnations of ISM involved computational reassignment^23,24^, with speeds limited by both acquisition and computation times, though made faster with the advent of the Zeiss AiryScan^25^. ISM can also be achieved all-optically^26,27^, and generalized to multi-point scanning^28,29^. Optical reassignment techniques have the advantage of directly providing superresolved images without the need for post-processing, however, they lack the versatility and flexibility of computational approaches. For example, they cannot on the fly compensate for differences between excitation and detection PSFs, or PSF distortions induced by local aberrations, etc.. More recently, the advent of SPAD array detectors has reignited an interest in computational ISM^30,31^. These are small arrays of detectors, typically 5×5 in size^32^, that respond at GHz rates, enabling much faster imaging than conventional camera-based approaches, even extending ISM to fluorescence lifetime imaging applications^30,31,33,34^. However, as small as a 5×5 array size appears, it imposes the acquisition of 25 signals through 25 independent electronic channels with high bandwidth and low noise, quickly making the electronics of ISM unwieldy.

We introduce an alternative approach to obtaining superresolution imaging while maintaining high light collection efficiency with less hardware complexity. The crux of our approach involves replacing the array detector of an ISM with a centroid detector which, at any given time, reports only the total intensity incident on the detector and its 2D centroid location. That is, it reports almost an order of magnitude reduced data compared to array detectors conventionally used for ISM. Our technique, called Centroid Reassignment Microscopy (CRM), is a simple variation on conventional confocal microscopy that is compatible with conventional (non-blinking) fluorophores. CRM combines the benefits of centroid localization, associated with localization-based superresolution approaches^11,12^, and pixel reassignment, associated with ISM. Compared to ISM, CRM provides resolution that surpasses the closed-pinhole limit (we demonstrate here resolution as fine as ∼100 nm), while being simpler to implement and less susceptible to detector noise. In addition, CRM automatically corrects for local aberrations and/or mismatches between excitation and detection PSF sizes without user intervention, enabling the possibility of extended depth of field (EDOF) imaging. In short, CRM is a uniquely attractive and versatile alternative to current superresolution approaches specifically designed for live sample imaging, as we demonstrate below.

## RESULTS

### Principle

CRM and ISM are similar in configuration (see Fig. 1a). A single laser-beam focus is scanned in two dimensions across a sample, producing a scanning excitation spot centered at ρ_ex_ = {*x*_ex_, *y*_ex_} with intensity defined by the excitation PSF. The resultant fluorescence generated at position ρ_em_ = {*x*_em_, *y*_em_} is then de-scanned through the same scanning mechanism and imaged onto an array detector with detection coordinate ρ_d_ = {*x*_d_, *y*_d_} and detection efficiency defined by the detection PSF. Because of de-scanning, ρ_d_ = ρ_em_ − ρ_ex_ and ρ_d_ = 0 corresponds to the location of the excitation spot at all times. If the fluorescence detected at position ρ_d_ is reassigned to the excitation spot location (ρ_d_ = 0), the array detector plays the role of a bucket detector (Fig. 1b top), and the resultant image becomes a conventional widefield image, yielding only modest resolution. The resolution can be enhanced by inserting a small pinhole at the excitation spot location (ρ_d_ = 0), producing a confocal image but at the cost of significant light loss. A similar resolution enhancement can be achieved by reassigning the fluorescence at ρ_d_ to a position intermediate between ρ_d_ and the excitation spot location (typically halfway for similar excitation and detection PSFs – see Fig. 1b middle). This is the principle of ISM, which has the advantage over confocal that it does not involve a pinhole and thus preserves full light collection efficiency. In CRM, we adopt yet an alternative approach where, for every scan position ρ_ex_, the detected fluorescence at position ρ_d_ is reassigned to its overall centroid location ρ_d_ (Fig. 1b bottom). This exploits the same advantage as ISM that it does not involve a pinhole, but, in addition, it does not require a full array detector such as shown in Fig. 1a,b. All that is needed is a centroid detector.

**Fig. 1.**
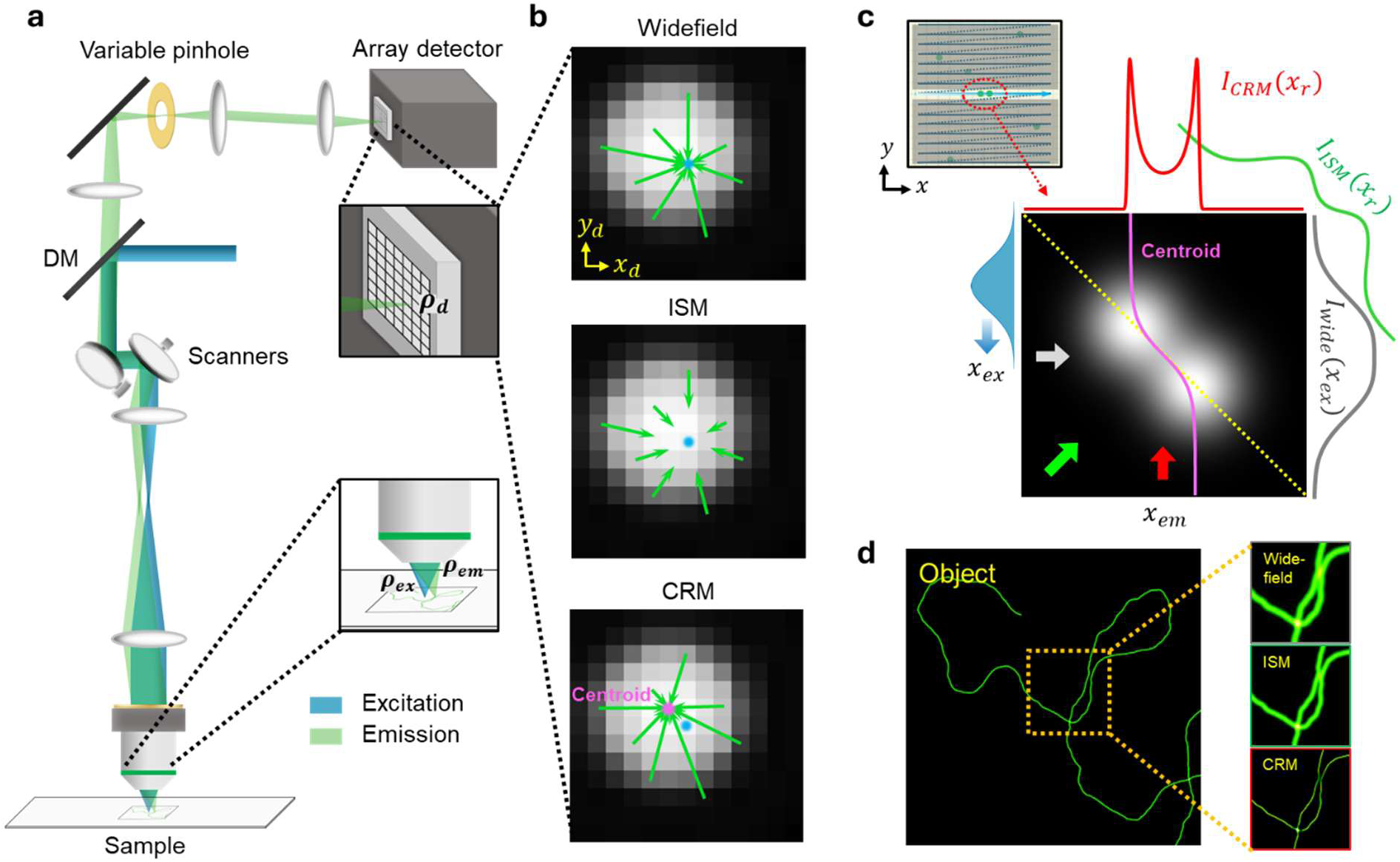
Principles of ISM and CRM. **a.** General schematic of scanning microscope. **b.** Example of emission distribution incident on array detector at a given scan time, where distribution happens to be slightly offset from location of excitation focus (ρ_d_ = 0, blue dot). **top:** In case of widefield microscopy, entire distribution (bucket detection) is reassigned to excitation focus location. **middle:** In case of ISM, distribution is pixel-wise reassigned roughly halfway to excitation focus location. **bottom:** In case of CRM, entire distribution is assigned to location of emission centroid (magenta dot). **c.** Simulation of *I*(*x*_ex_, *x*_em_) obtained from a single line of a 2D raster scan spanning pair of fluorophores located at same *y* coordinate (inset top left). In this example, excitation and emission PSFs are assumed Gaussian of equal size (blue-filled Gaussian indicates scanning excitation), and fluorophore separation is half the PSF width. Reassignment operations depicted in **b** correspond to projection operations depicted in **c**. Widefield image (gray trace) is obtained from projection of *I*(*x*_ex_, *x*_em_) along *x*_em_ (gray arrow). Fluorophores are insufficiently spatially separated here to be distinguished by widefield microscopy. ISM image (green trace) is obtained from projection along anti-diagonal (green arrow). Because fluorophores appear sparser along this projection direction, they become distinguishable. CRM image (red trace) is obtained from projection of weighted centroid trajectory (magenta curve) along *x*_ex_ (red arrow). Because of temporal sparsity induced by scanning, the centroid trajectory follows a nonlinear path, enabling fluorophores to be readily distinguished by CRM. *x*_r_ is reassignment coordinate. **d.** Schematic of resulting images of fibrillar object.

The origin of resolution gain in CRM is somewhat different than in ISM. In ISM, information about the origin of detected photons is derived equally from detection and excitation PSFs, and the resultant spatial map of fluorescent emitters becomes accurate only after scanning. CRM, instead, is based on the principle of localization, where information about the location of fluorescent emitters comes dominantly from the detection PSF. For such localization to be accurate, a condition of emitter sparsity must be met. In conventional localization microscopes based on uniform illumination, emitters are rendered sparse in time by the use of blinking fluorophores. More recently, localization microscopes have exploited the gain in sparsity that comes from the use of time-varying structured illumination^17,18^ which improves localization accuracy. In CRM, we exploit this same gain in sparsity that comes from time-varying structured illumination, but without the imposition of blinking fluorophores. Because the excitation focus is scanned, emitters appear distributed not only in space but also in time. This is illustrated in Fig. 1c, depicting two emitters in close proximity. In the case of widefield microscopy, the spatial separation between the emitters is not sufficient for them to be distinguished. In the case of CRM, the centroid of the fluorescence produced by the emitters follows a sigmoidal trajectory (magenta trace) as a function of scan position ρ_ex_ that jogs from one emitter to the other as a function of time. Because of its nonlinear shape, the trajectory spends more time centered on the emitter locations than elsewhere, thus allowing more signal to accumulate at the emitter locations than elsewhere. As a result, the gap between the two emitters becomes readily apparent in the final reassigned image (Fig. 1c), and resolution becomes enhanced. The degree of enhancement depends on a variety of factors, such as the possible presence of a pinhole (for purposes of optical sectioning) and also the distribution of the emitters themselves. In general, the sparser the distribution of emitters, the greater the enhancement. However, as evidenced by theory and simulation (Supplementary Notes 1-7) and by experimental results (see below), CRM is shown to enable finer resolution than ISM even for relatively dense distributions of emitters.

### CRM image formation

The only hardware differences between a CRM and a conventional confocal microscope are that we make use of a largely opened pinhole and a centroid detector. Different types of centroid detectors are available. Here, to achieve sufficient bandwidth, we make use of a simple quadrant detector (QD – Fig. 2a). However, by itself, a QD cannot report centroid location. An additional scaling factor is needed related to the spot size incident on the QD. This scaling factor can be estimated a priori based on the known magnification and resolution of the optics. Alternatively, it can be measured experimentally using a method of image cross-correlation 35,^30^.

**Fig. 2.**
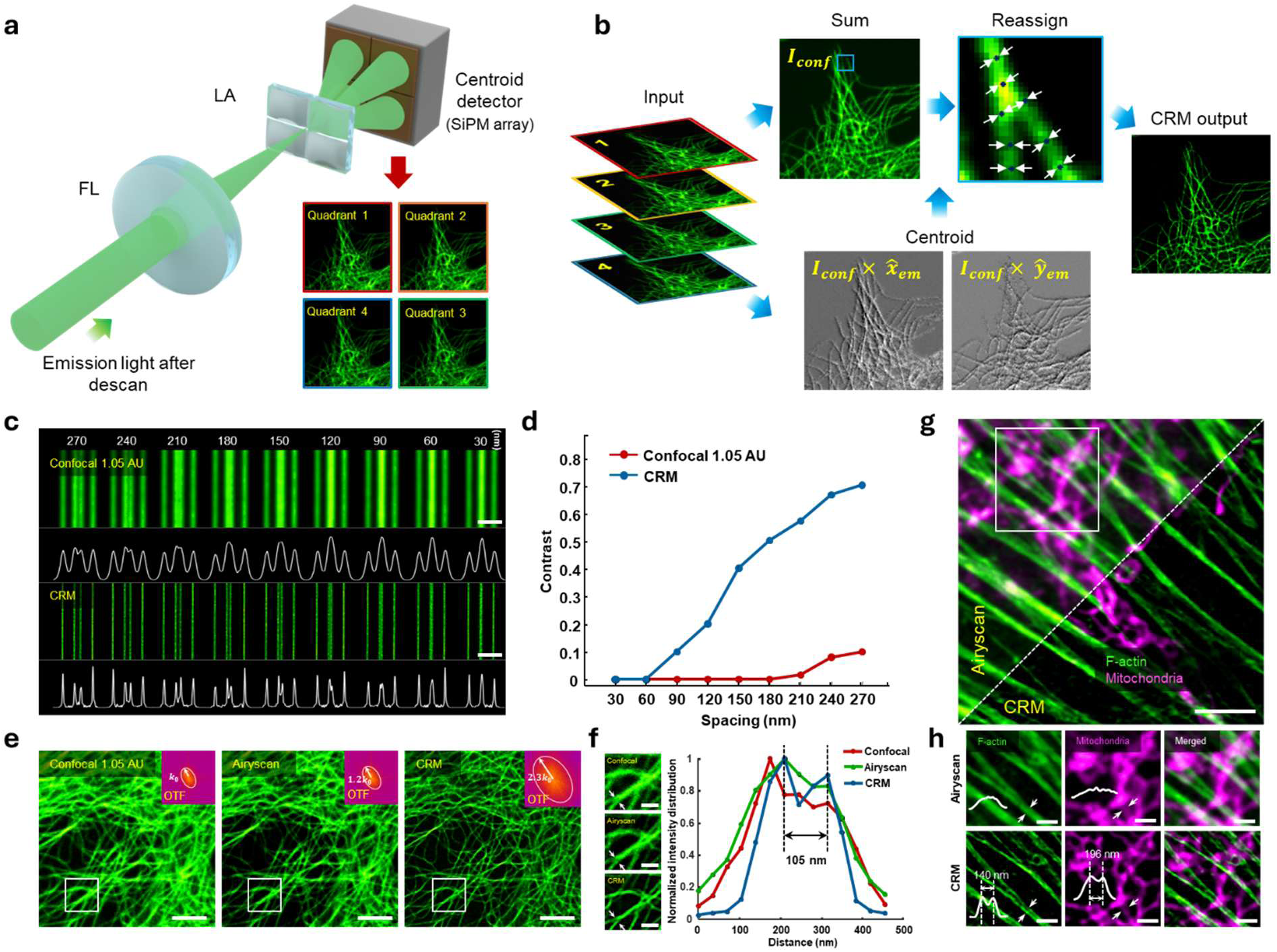
CRM image formation with a centroid detector. **a.** Emission light through a moderately open pinhole (1.05 AU, not shown) is focused onto a lenslet array (LA) and detected by a quadrant SiPM array. The quadrants obtain 4 raw images upon scanning, which serve as inputs to the CRM processing pipeline. **b.** For each excitation scan position ρ_ex_, the emission total intensity *I*_conf_ and centroid location 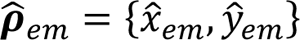 are calculated from the raw images and used to perform CRM (more details in Methods). **c.** Resolution target (Argo-SIM V2) imaged by confocal (summed quadrants, pinhole 1.05 AU) and CRM. Scale bar: 1 μm. **d.** Plots of contrast of central line pairs obtained from **c** enable estimate of CRM resolution (here ∼90 nm, for 488 nm excitation with a 1.42 NA objective). **e.** Anti-α-tubulin labeled microtubules in the fixed BPAE cell imaged by confocal (left) and CRM (right), compared with ISM image of the same region (middle -- obtained with Zeiss LSM 980 Airyscan2, pinhole 1.25 AU). Insets depict spatial frequencies (log scale). Scale bar: 2 μm. **f.** Zoomed areas from white boxes in **e**, and line profiles across spans indicated by white arrows. Scale bar: 600 nm. **g.** Two-color images (obtained sequentially) of Alexa Fluor 488 phalloidin-labeled F-actin and MitoTracker Red CMXRos-labeled mitochondria in fixed BPAE cell imaged by CRM and Airyscan. Scale bar: 3 µm. **h.** Zoomed areas from **g** with intensity profiles across spans indicated by the white arrows. Scale bar: 800 nm. Experiments were repeated three times independently with similar results.

In Fig. 2b, we provide a schematic of our CRM algorithm based on the use of a QD (more details provided in Methods, and Supplementary Fig. 8). As noted above, CRM is generally operated in the presence of a moderately open pinhole, here typically 1.05 Airy units (AU) in size. From the raw images produced by the four detector quadrants, we thus derive a conventional confocal image corresponding to the summed four images, as well as a 2D centroid map characterizing the detected centroid trajectory as a function scan position ρ_ex_. To increase robustness to noise, we apply a lowpass filter to the centroid map with a tunable kernel size typically set to no larger than half the widefield PSF, roughly corresponding to the filtering resulting from the doubled spatial frequency support of an ideal confocal of zero-size pinhole. The resolution gain provided by CRM can be quantified using a calibrated resolution target (Argo-SIM V2) that serves as a ground truth. Figure 2c compares confocal and CRM images of line pairs of decreasing separation. With CRM, line-pairs as close as 90 nm can be readily distinguished (Fig. 2d, contrast calculation in Methods), providing an estimate of CRM resolution (with 488nm excitation and 1.42 NA oil-immersion objective). We further compare the resolution gains provided by CRM and ISM, making use of a state-of-the-art commercial ISM (Zeiss LSM980 Airyscan2 -- see Methods). Figure 2e shows images of fixed anti-α-tubulin labeled microtubules in fixed Bovine Pulmonary Artery Endothelial (BPAE) cells obtained with confocal (left, summed quadrants) and CRM (right), compared with the same region obtained with ISM (middle). For proper comparison, the illumination powers at the sample, pixel sizes and pixel dwell times were set to be equal for all modalities. CRM provides improved resolution of microtubule separation, as shown in Fig. 2f. We further performed sequential two-color imaging of BPAE cells (Figs. 2g,h: F-actin in green, mitochondria in magenta), again illustrating the improved separation of features enabled by CRM. We note that the resolution improvements obtained with both CRM and ISM depend on their respective degrees of reassignment (parametrized by a reassignment factor α – see Supplementary Notes 1,4,7 and Extended Data Figs. 2,3). All comparisons of CRM with ISM throughout our main text are obtained with α chosen to provide optimal resolution (α → 1 for CRM; α ≈ 0.5 for ISM). We further note that the non-zero size of the individual detector elements in ISM (here 0.2 AU) can limit the overall ISM resolution, whereas CRM is not subject to this limitation.

### CRM resolution improvement is robust to noise

Major advantages of CRM are its high light collection efficiency and improved resolution capacity compared to ISM for equal photon budget. In Fig. 3, we provide an overview of the effect of different illumination powers (hence different SNR levels) on image resolution. Note that from the same Airyscan raw data we can synthesize a standard confocal image (summation of all detector pixels, 1.25 AU pinhole size), a small-pinhole confocal image (central detector pixel only, 0.2 AU pinhole size), and an ISM image (reassigned detector pixels). These are compared with images obtained from our CRM, again with equal illumination powers at the sample, pixel size and pixel dwell times. A now standard method of evaluating microscope resolution, particularly suitable for superresolution imaging, is the method of Fourier Ring Correlation (FRC) ^36^, which has the advantage that it takes into account the detrimental effects of detection noise (shot noise, electronic noise, etc.) on resolution. Again, CRM is observed to provide the highest resolution (Fig. 3c). Because CRM provides improved resolution compared to ISM for equal photon budget, the detected photons become compressed into smaller areas, leading to improved SNR (Supplementary Fig. 9). The resolution enhancement provided by CRM becomes all the more significant at lower excitation powers where a reduced SNR more severely undermines image quality (particularly apparent in the case of the small-pinhole confocal).

**Fig. 3.**
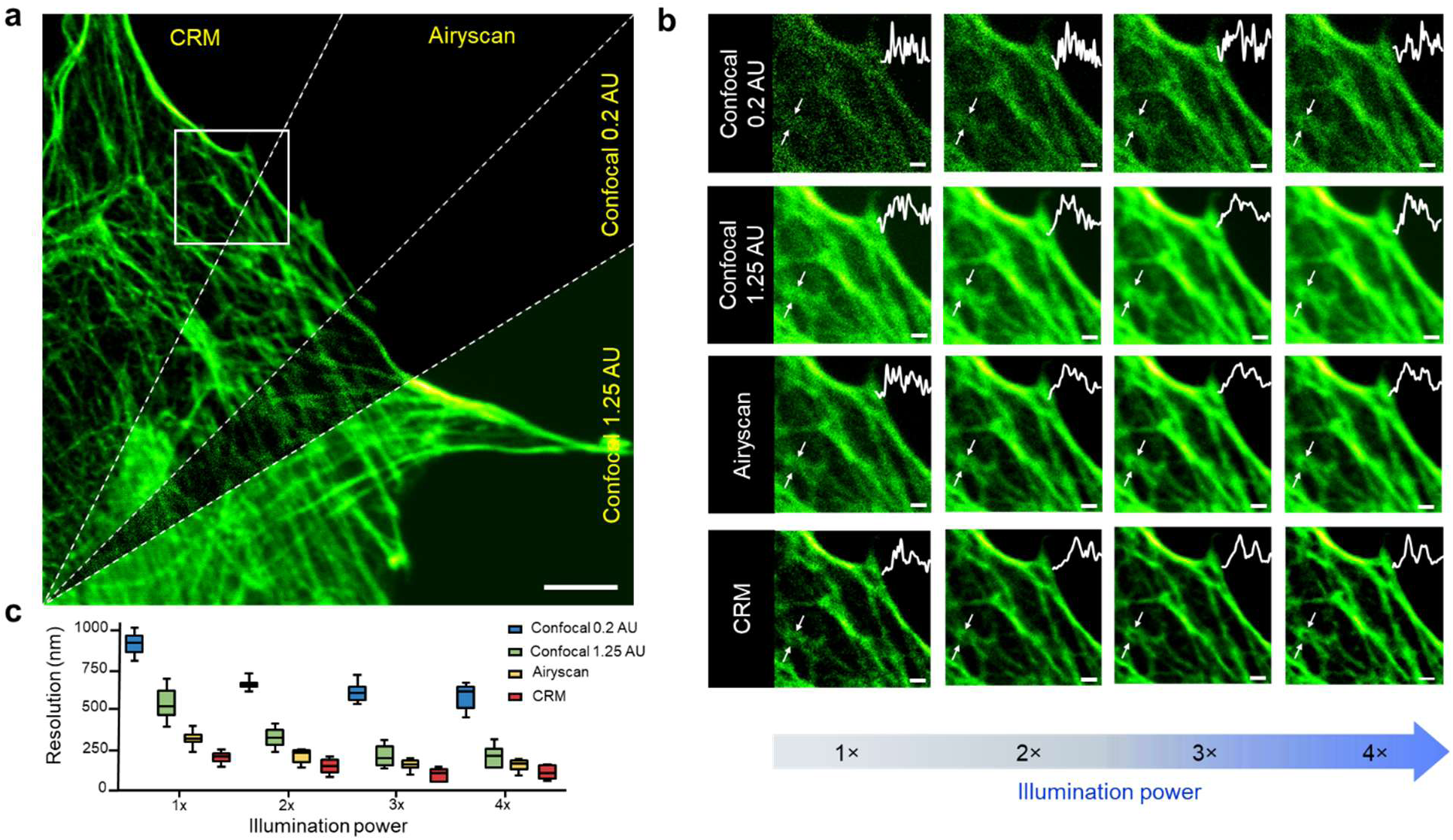
Demonstration of CRM resolution enhancement and robustness to noise. **a.** Comparison of images of BPAE F-actin labeled with Alexa 488 phalloidin obtained from CRM and Airyscan. Counterclockwise from bottom: Airyscan sum mode (confocal 1.25 AU), Airyscan center channel only (confocal 0.2 AU), Airyscan SR mode (without deconvolution), and CRM (pinhole 1.05 AU). Scale bar: 3 µm. **b.** Zoomed area from the white box in **a** obtained with increasing illumination power (1× corresponds to 1.25 μW at sample). **c.** Spatial resolution inferred from FRC analysis of F-actin images (FRC threshold used to define resolution set to 1/7, as per convention; experiments repeated over 8 distinct ROIs). Spatial resolutions for 4× power given by: CRM: 107 nm ± 35 nm; Airyscan: 160 nm ± 35 nm; Confocal 1.25 AU:215 nm ± 59 nm, and Confocal 0.2 AU: 588 nm ± 73 nm). Experiments were repeated three times independently with similar results.

### Live sample imaging with CRM

CRM is highly light-efficient, making it well suited for long-term imaging of live samples. In Fig. 4, we provide examples of live-sample imaging of dynamic processes such as mitochondrial fusion and fission in U2OS osteosarcoma cells and neuronal motion in quasi-freely moving jellyfish. For the U2OS experiments (Fig. 4a-d), we recorded a 10-minute time-lapse video of a single U2OS cell using gated illumination (see Methods). CRM is able to resolve morphological dynamics in U2OS mitochondria more effectively than standard confocal microscopy (summation of four quadrants), enabling us to more clearly observe mitochondrial fusion (Fig. 4b and Supplementary Video 1) and fission events (Extended Data Fig. 4 and Supplementary Video 1) over time scales consistent with previous reports^1,2^. This trend is more apparent in the 1-hour time-lapse sequence (Fig. 4c-d, and Supplementary Video 2) obtained under ambient conditions with lower intensity continuous illumination (see Methods and Supplementary Table 2). CRM delivers both robust spatial resolution improvement and increased signal-to-background ratio (SBR) (Fig. 4d). The reason for SBR improvement is that signal structure becomes reassigned by CRM into smaller areas, while background, which is mostly uniform, produces no centroid displacement and thus remains largely untouched. Note that the integrated preamplifier in our QD limited our maximum pixel rate to about 2 MHz, which nevertheless allowed us to attain video-rate (30 Hz) imaging for frames as large as 256×256 pixels (Extended Data Fig. 5 and Supplementary Video 3). For live jellyfish imaging, we also used continuous illumination. As is apparent from the extracted frames (Fig. 4e-f) and full videos (Supplementary Video 4 & 5), CRM enables much finer features to be resolved in the motion dynamics of two neuronal bodies, again with increased SBR.

**Fig. 4.**
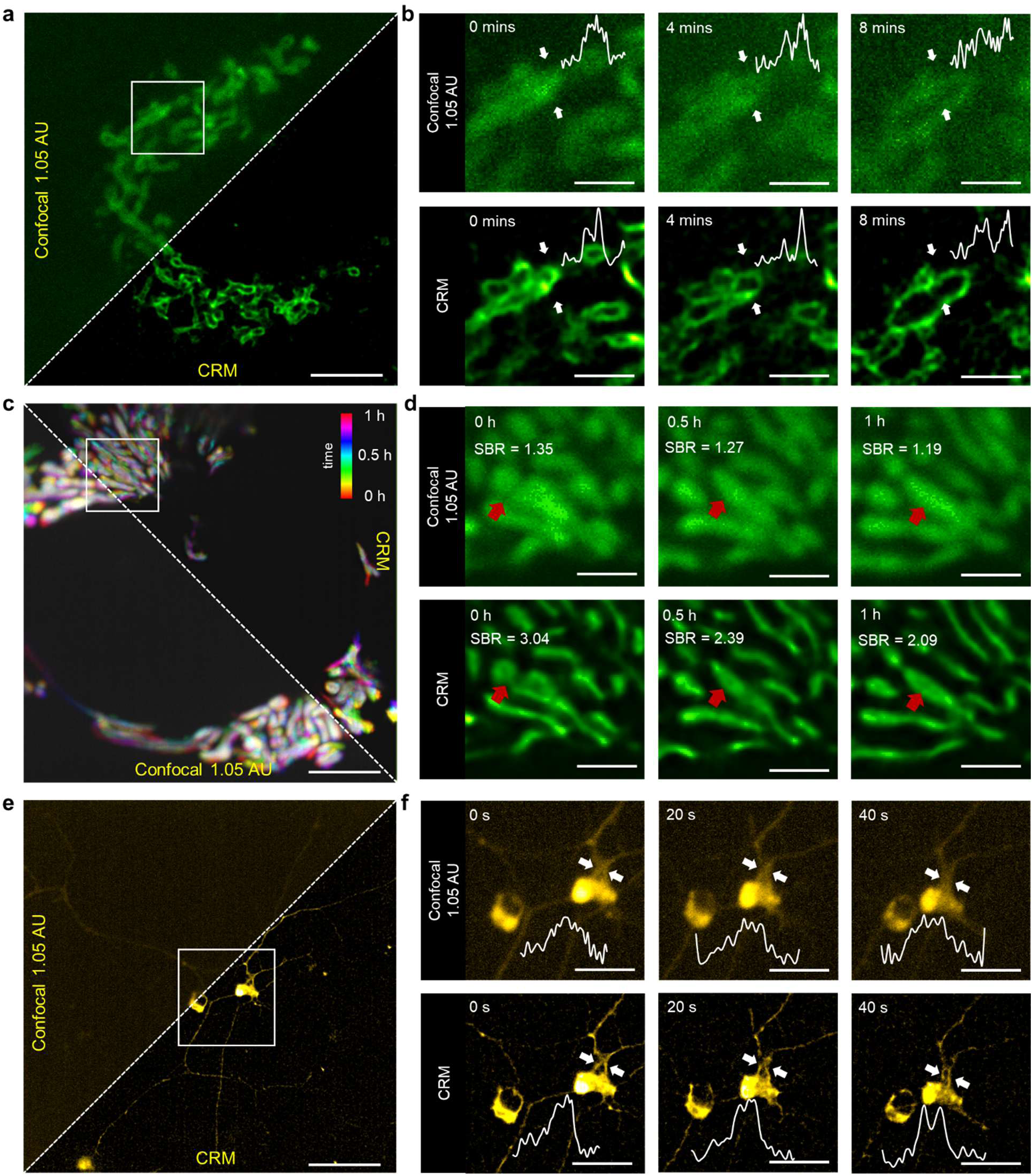
Demonstration of live-sample imaging with CRM. **a.** Mitochondrial dynamics in live U2OS osteosarcoma cell expressing an EGFP-tagged mitochondrial outer membrane protein (TOM20-EGFP) captured by 60×, 1.42NA objective. FOV: 16 µm × 16 µm. Scale bar: 3 µm. Full video shown in Supplementary Video 1. Conventional confocal image (pinhole 1.05 AU) obtained by summing of raw quadrant images. **b.** Time lapse of mitochondrial fusion from zoomed region (white box in **a**). Normalized intensity profiles across spans indicated by white arrows. Scale bar: 1 µm. Full video shown in Supplementary Video 1. **c.** Temporal projection across 1 hour of mitochondrial network activity in live U2OS osteosarcoma cell (same in **a**). FOV: 22 µm × 22 µm. Scale bar: 3 µm. Full video shown in Supplementary Video 2. **d.** Time lapse of mitochondrial interactions from zoomed region (white box in **c**). The red arrow highlights a shape change upon interaction between mitochondria. Scale bar: 1.5µm. Full video shown in Supplementary Video 2. **e.** mCherry-expressing RFamide neuron dynamics in live jellyfish (*Clytia hemisphaerica*) captured by 20×, 1.0 NA objective. FoV: 49µm × 49µm. Scale bar: 11 µm. Full video is shown in Supplementary Video 4. **f.** Time-lapse of neuron bodies and neural processes from zoomed region (white box in **c**). Normalized intensity profiles across spans indicated by white arrows. Scale bar: 4 µm. Full video is shown in Supplementary Video 5. Similar imaging experiments were performed three times with similar results.

### CRM enables EDOF imaging

In practice, it is often preferable to operate CRM with a pinhole that is somewhat closed so as to benefit from a degree of optical sectioning dependent on pinhole size. In the event that the pinhole is largely open, the optical sectioning becomes weak, meaning that out-of-focus signal becomes incident on the QD. In effect, this leads to a broadening of the detection PSF, and hence to an increase in the scaling factor *s*→ required for centroid calculation. Inasmuch as our estimation of *s*→ is based on quadrant image cross-correlation (see Methods), CRM is able to adaptively adjust for such PSF broadening. Moreover, since the estimations are performed locally, the reassignment scaling factors can vary across the sample. If parts of the sample are more out of focus than others, CRM adaptively adjusts for this. As a result, CRM enables clear visualization of fine structures even when the sample exhibits depth variations. That is, CRM is robust to defocus aberrations, enabling the possibility of extended depth of field (EDOF) imaging. To demonstrate this, we imaged a 3D staircase of cylinders in an Argo-SIM slide which served as a ground-truth sample (Fig. 5a-b). Similar results are obtained with jellyfish imaging, shown in Fig. 5c, where the out-of-focus blur apparent in the confocal images (left) is largely compensated by CRM (right) across a depth range of 10 μm. The latter can be compared with a confocal stack obtained with a 0.5 AU pinhole (Fig. 5d and Extended Data Fig. 6). Though not demonstrated here, CRM is expected to be robust to more general aberrations than defocus, provided these do not cause an overall shifting of the centroid of the detection PSF.

**Fig. 5.**
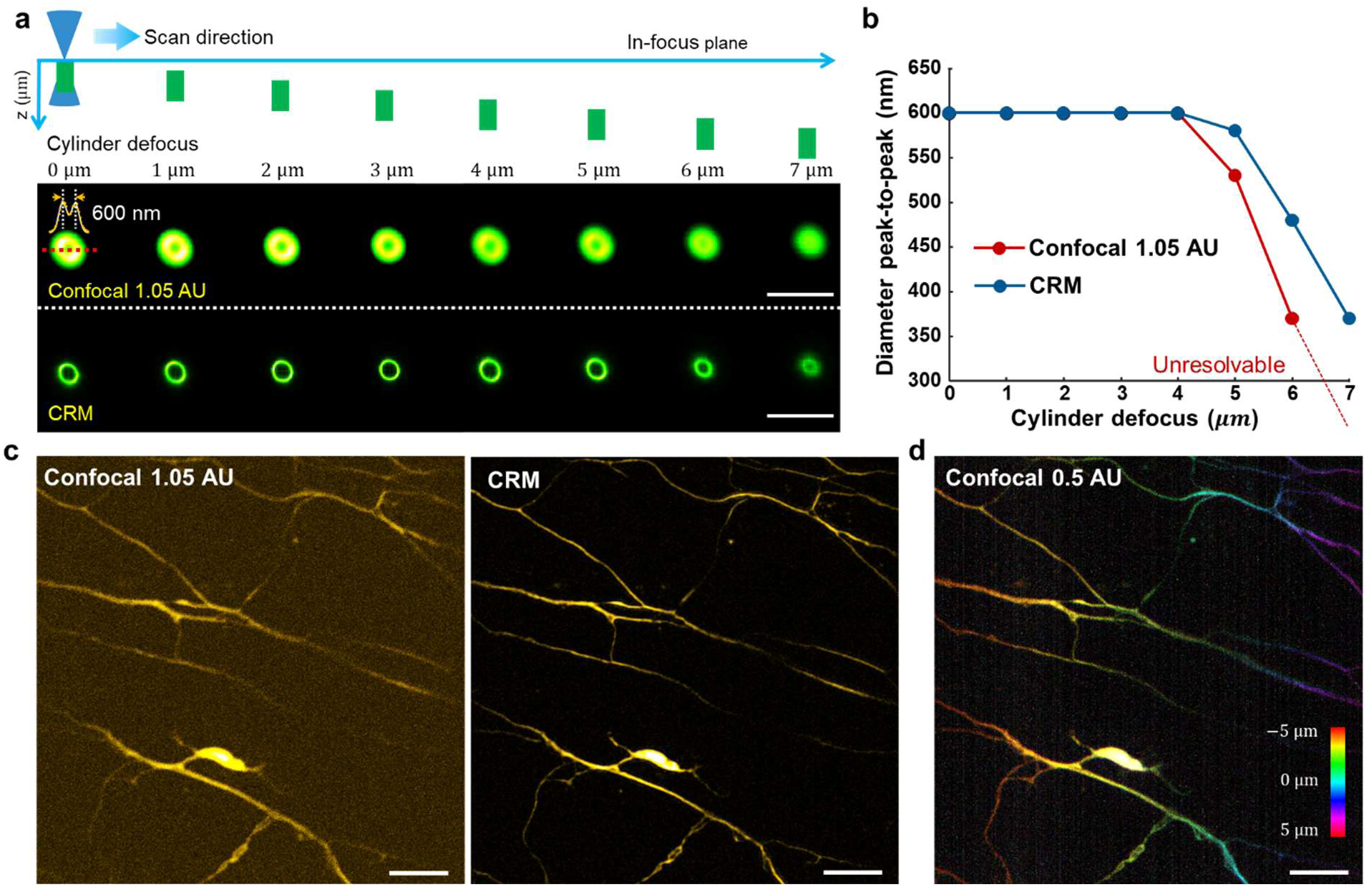
CRM provides virtual EDOF imaging. **a.** Confocal (summed quadrants; pinhole 1.05 AU) and CRM images of calibration targets (Argo-SIM V2) consisting of empty cylinders of progressively increasing defocus distance (1 µm step). Excitation scan plane is focused at top of left-most cylinder. CRM consistently provides better resolution than confocal. Scale bar: 3 µm. **b.** Measured peak-to-peak diameters of cylinders (600 nm, inferred from a double Gaussian fit) are better preserved over a larger range of defocus distances. **c.** Neurons labelled with mCherry in live jellyfish captured by confocal (left) and CRM (right). Scale bar: 3 μm. **d**. Projection of an image stack of the same region in **c** obtained with a custom-built standard confocal microscope (pinhole 0.5 AU) with a z-step of 1 μm across a depth of 10 μm. Color corresponds to depth. Scale bar: 3 μm. Similar imaging experiments were performed three times with similar results.

## DISCUSSION

Different metrics can be used to characterize the resolution of a microscope^36^, in either the Fourier domain (e.g. the Abbe limit) or the pixel domain (e.g. the Rayleigh limit). The former is often readily apparent when viewing the intensity spectra of images, as illustrated in Fig. 2e (insets). The latter, which characterizes the separability of two emitters, is generally more relevant for biological imaging since organelles of interest tend to be localized. We have shown that CRM provides better emitter separability than both confocal microscopy and ISM. Compared to a confocal microscope, CRM is similarly simple and can be implemented as an add-on with minimal modifications to any conventional confocal microscope, provided the latter is equipped with 4-channel electronics. The main benefit of CRM is that it maintains full detection efficiency since it does not require the use of a closed pinhole. This benefit preserves SNR even when imaging at low light levels, such as encountered when imaging at high speed or with reduced excitation powers, as often imposed with live-sample imaging. Compared to ISM, CRM makes use of fewer detectors, simplifying the detection electronics and mitigating electronic noise, all while providing sharper images for equal photon budget.

However, CRM does not come without drawbacks. The main drawback is that it is nonlinear, meaning that the CRM imaging process cannot be described as a convolution with a PSF. On the one hand, it is precisely this nonlinearity that facilitates emitter separability and leads to sharper images than ISM. On the other hand, when image nonlinearity is a concern, this can be mitigated by various image regularization approaches (Supplementary Note 6). Another drawback of CRM is that, because it avoids the use of a pinhole, it sacrifices the main benefit that comes from a pinhole, namely, a capacity for optical sectioning by physical rejection of out-of-focus background. In practice, we have found that CRM works best in combination with a pinhole that is not fully opened (similar to ISM), thus benefiting from some degree of optical sectioning without undue sacrifice of detection efficiency. The presence of a pinhole has the additional effect of smoothing the CRM images. Of course, there may be scenarios where a weak optical sectioning capacity is actually desired, for example, to mitigate artifacts that can arise from axial sample motion. CRM maintains high resolution in this case, even over an extended focus range, since it is inherently robust against defocus aberrations.

In recent years, there have been reports of single- or few-frame enhanced-resolution imaging of conventional fluorophores based on the use image post-processing^37–39^ or machine learning^40,41^. We emphasize that CRM, in its current form, does not rely on a priori knowledge about the sample, though it may benefit from the aid of such techniques in future implementations. Additionally, one could envisage reducing the number of detection channels either by using 3-element detectors or true non-pixelated centroid detectors, further streamlining the CRM hardware. In conclusion, because of its simplicity and broad applicability, we believe CRM will be of general interest to the biological imaging community.

## METHODS

### Imaging methods

#### CRM setup

A detailed schematic of our CRM setup is shown in Supplementary Fig. 5, with its components listed in Supplementary Table 1. Essentially, CRM is the same as a conventional point-scanning confocal microscope, except that the pinhole is opened and the single-element detector is replaced with a centroid detector. In the illumination path, two lasers (Omicron LuxX® 488-200 and Vortran Stradus®561-50) were combined by a dichromatic mirror (DM1, Chroma ZT488/561rpc). To obtain single-mode beam output, spatial filters (PH1, L1, PH2, and L2) were placed in front of the lasers. A point focus was imaged into the sample plane by a series of relay lenses (L2, PL1, PL2, SL, and L3) and an objective (Olympus XLUMPLFLN 20× /1.0 NA W or UPlanXApo 60×/1.42 NA Oil). The focus was scanned across the field of view (FoV) by galvanometric scanners GS-X, and GS-Y (ScannerMAX Saturn 5B). To reduce FoV-related aberrations, we separated the galvanometric mirrors by a unit-magnification 4f relay system comprising two Plössl lenses (PL1 and PL2)^42^. achieving a uniform FoV of −5° to + 5° over a spectral range 488 nm to 620 nm, as verified by a Zemax OpticStudio simulation (details provided in Supplementary Fig. 6). A 5× magnification relay consisting of a scan lens (SL) and a doublet lens (L3) was inserted between GS-Y and the objective pupil plane to ensure full coverage of the objective back aperture.

Emitted fluorescence was epi-collected, de-scanned, and separated from the reflected illumination light with a dichromatic mirror (DM2, Semrock Di01-R488/561). The de-scanned fluorescence was then focused onto a pinhole wheel (PW, PHWM16, Thorlabs) by a doublet lens (L4), and the pinhole size could be easily changed by rotating the wheel, providing different optical sectioning capacities. Two pinholes of diameters 150 μm (for 60×/1.42 NA objective) and 70 μm (for 20×/1.0 NA objective) were typically chosen based on the different estimated lateral magnifications, 333× (for 60×/1.42 NA) and 111× (for 20×/1.0 NA). To minimize crosstalk from the illumination light, an additional notch filter (Thorlabs NF488-15 or NF561-18) was inserted in the detection path and mounted on a filter wheel (FW). Sequential two-color imaging was enabled by software-controlled toggling of the respective lasers and rotation of the FW. The image of the pinhole was then relayed by two doublet lenses (L5 and L6) and delivered to the centroid detection module. Samples were mounted on a motorized two-axis translation stage (Thorlabs PLSXY) and controlled by a stepper motor controller (Thorlabs MCM3000).

#### Centroid detector design

Different from a conventional point-scanning confocal microscope, a CRM makes use of a centroid detector rather than a single-element detector. Quadrant detectors (QDs) are often used as centroid detectors^43,44^, however, these can suffer from reduced fill factor owing to the gaps between detector elements. To improve fill factor, the fluorescence focus was first projected onto four adjacent lens corners in a lenslet array (LA, the four central lenslets of a Thorlabs MLA1M1), with design considerations shown in Supplementary Fig. 7. The resultant four quadrants of the fluorescence focus were then collected by a 2 × 2 SiPM array (the four central elements in a Hamamatsu C13369-3050EA-04 module), thus steering the fluorescence from the gaps between the detector elements.

#### Conventional point-scanning confocal microscope

To enable comparison of CRM with conventional point-scanning confocal microscopy, a single-element detection path was also included in our setup. A flip mirror was introduced after the pinhole-image relay, which routed the fluorescence to a single-element SiPM module (Hamamatsu C13366-3050GA). By switching pinhole size and flipping the mirror in and out of the detection path, the system could be switched from conventional confocal microscopy to CRM.

#### Airyscan microscope

To enable comparison of CRM with ISM, we made use of a state-of-the-art commercial Airyscan microscope (Zeiss LSM 980 Airyscan2)^25,45^. All experiments performed with Airyscan were made using a same objective (Zeiss Plan-Apochromat 63×/1.4 Oil DIC M27). When comparing CRM and ISM imaging, all settings such as laser power at the sample, frame rate, pixel size, pixel number, etc., were adjusted to be the same.

### Calibration

Pixel size was calibrated before each experiment. To determine the pixel size, a fixed 4 μm bead slide (Invitrogen T14792 TetraSpeck™ Fluorescent Microspheres Size Kit) was translated by a known distance, *d*_calibration_. Two images were acquired before and after translation. The number of pixels the beads shifted, *N*_pixels_, was then used to calculate pixel size as 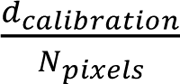.

Illumination power was measured prior to all experiments (see Supplementary Table 2). To determine the power level at the sample plane, we used a power sensor (Thorlabs S130C) to measure the focused beam after passing through the objective, oil or water media, and cover glass. Readings were recorded using a power meter (Thorlabs PM100D).

### Data acquisition system

CRM microscope control and timing were performed using ScanImage software (Vidrio SI 5.7 R1)^46^ running on MATLAB 2014b. Communication between the computer and scanner driver boards was performed with a PXIe-6341 multifunction I/O module (National Instruments). Analog signals from the SiPM array module were converted by a NI 5734 four-channel digitizer followed by a NI PXI-7961R FPGA module, all mounted in a PXIe-1073 chassis (National Instruments). For gated illumination, both laser sources were triggered by a USB-6356 DAQ (National Instruments), ensuring synchronization with the SiPM array. For z-stack acquisition, the objective lens was mounted on a motorized focusing module (Thorlabs PLSZ) driven by a compact stepper motor controller, with ScanImage generating the control commands for axial scanning. Experimental settings are listed in Supplementary Table 2.

### Centroid measurement with a quadrant detector

We employed a QD as a centroid detector. The four quadrants of the QD produce four sub-images, denoted by *I*_1,2,3,4_(ρ_ex_), such that 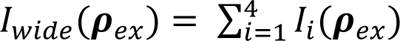 in the absence of a pinhole, or 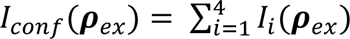 in the presence of a largely open pinhole (the notation *I*_confO_(ρ_ex_) is reserved for the case of a closed pinhole). However, a QD is an imperfect centroid detector since it cannot, by itself, provide centroid location in units of distance. An additional scaling vector is required. The centroid displacement vector 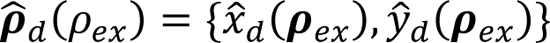 (location of centroid relative to the detector center) is thus given by,

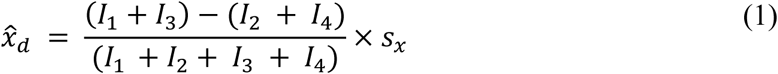

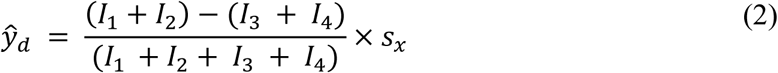

where *s*_x_ and *s*_y_ are the scaling factors, related to the spot size incident on the detector. We describe below how these scaling factors are estimated.

Equations (1) & (2) are linear approximations that are valid when the displacements of the emission spot from the center of the QD are smaller than the spot size itself. For larger displacements, the relation between QD response and actual spot displacement becomes nonlinear and saturates^43^. Throughout this work, we assume the displacements are small and the linear approximations are generally valid, bearing in mind that CRM is tolerant to errors, since inaccuracies in the estimations of displacements are tantamount to small errors in *α*. As shown in Extended Data Figs. 2,3 and Supplementary Figs. 1,4, variations in *α* lead simply to different degrees of sharpness in our final CRM image reconstructions. In other words, resolution enhancement remains preserved over a wide range of estimation errors.

### Estimation of scaling factors for centroid calculation

The simplest approach to estimating the scaling factors *s*_x_ and *s*_y_ is to assume they are given by the average size (FWHM) of the system PSF, which can be measured experimentally using a calibrated point target (Extended Data Fig. 1) or estimated theoretically based on a model of the microscope system. In practice, this assumption generally leads to quite acceptable results.

A more refined approach measures these scaling factors adaptively. Each quadrant of the QD produces a quasi-confocal sub-image given by

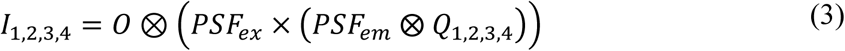

where *O* is the object, *Q*_1,2,3,4_ are the masks defined by the QD detector elements (further restricted by the pinhole, if present), and ⊕ indicates a convolution operation. Because *Q*_1,2,3,4_ are displaced from one another, so too are the sub-images *I*_1,2,3,4_, with a displacement distance corresponding to the system PSF size. By computationally registering the four images so that they are superposed, one can estimate *s*_x_ and *s*_y_ from the corresponding registration distances. This technique was described and demonstrated by simulation in Refs.^35^, where image registration was performed by simple image cross-correlation and shown to be equivalent to ISM pixel reassignment^30,31,34^. An advantage of this cross-correlation approach is that it requires no a priori knowledge of *PSF*_em_ and *PSF*_ex_ and automatically corrects for possible differences in their sizes (generally present because of the Stokes shift between emission and excitation wavelengths). In Refs.^30,31,34,35^, cross-correlations were performed globally over the full sub-images, leading to registration distances that were averaged across the entire FoV. This did not make allowances for possible local variations in *PSF*_em_or *PSF*_ex_ because of spatially varying aberrations. To take such aberrations into account, we subdivided our sub-images into zones and calculated local image cross-correlations, enabling us to infer the scaling factors *s*_x_ and *s*_y_ locally as opposed to globally. Smooth transitions from zone to zone were ensured by spatially lowpass filtering *s*→.

### Reconstruction algorithm

A detailed reconstruction pipeline is shown in Supplementary Fig. 8. After acquiring the four sub-images from each quadrant, we typically applied optional background subtraction using a local minimum filter^39^. The sliding window size for the local filter was typically set to match the largest structural feature in the image. For fair comparison, Airyscan images were also subjected to the same operation with the same parameters before the pixel reassignment. Because of the normalization (division) operations in Eqs. (1) & (2), noise can be amplified at low signal levels, possibly resulting in errors in the centroid estimation. To suppress such errors, a Gaussian lowpass filter was applied to smooth the centroid maps. The filter kernel size was chosen to be slightly smaller than the effective system PSF size, leading to a tradeoff between resolution enhancement and noise suppression. At each image pixel, the emission centroid location was then calculated from 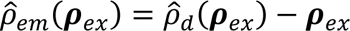, and CRM image reconstruction was performed according to Eq. (S3) in Supplementary Note 1, making use of a user-defined reassignment factor α (generally set to 1).

For ISM image reconstruction, we followed the pixel reassignment procedure described in Ref. ^24^, taking into account the difference between excitation and emission wavelengths when presenting optimized ISM results.

### Data processing and analysis

Data processing was performed using custom code written in MATLAB 2024b. The computational environment included an AMD Ryzen 5 7600X3D 6-core processor (4.1 GHz, 12 logical threads) and 32 GB RAM. Processing parameters are provided in Supplementary Table 3.

To quantify the resolution enhancement achieved by CRM, particularly under noisy conditions, FRC analysis was performed. FRC requires two statistically independent images of the same scene under identical imaging conditions^36^. In our study, we captured the same sample region with two consecutive exposures. FRC calculations were conducted using custom MATLAB scripts, following the methodology described in Ref. ^36^.

ROI line profiling and data extraction were carried out in Fiji. Corresponding intensity profiles and statistical analyses, including FRC resolution plots, were generated using Origin 2018.

For quantitative contrast analysis, we used

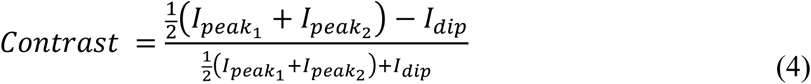

To estimate the cylinder diameter from the peak-to-peak distance, we fitted the line profile through the diameter with a double-Gaussian function using the MATLAB Curve Fitting Toolbox.

### Sample preparation

#### Argolight calibration slide

We used the Argo-SIM V2 slide (Argolight), and imaged two patterns. The first is the resolution target of gradually spaced lines imaged using a 60×/1.42NA objective. The second is a 3D crossing stair of cylinders with axial step size 1 µm imaged using a 20×/1.0NA objective.

#### Fixed sample slides

Fluorescent beads and fixed BPAE cells slides were purchased from Thermo Fisher Scientific.

Beads: T14792, Invitrogen™ TetraSpeck™ Fluorescent Microspheres Size Kit. This kit contains one microscope slide with six viewing regions, five of which contain microspheres of size 0.1, 0.2, 0.5, 1.0 or 4.0 μm respectively.

BPAE cells: FluoCells™ Prepared Slide #1 (BPAE cells with MitoTracker™ Red CMXRos, Alexa Fluor™ 488 Phalloidin, and DAPI) and FluoCells™ Prepared Slide #2 (BPAE cells with Mouse Anti-α-tubulin, AlexaFluor™ 488, FL Goat Anti-Mouse IgG, Texas Red™-X Phalloidin, and DAPI).

#### Stable live U2OS cell line generation and preparation

U2OS cells (Sigma-Aldrich, 92022711-1VL) were maintained at 37 °C and with 5% CO2 in culture media consisting of Dulbecco’s Modified Eagle Medium (Cytiva, SH30285.01), containing 10% fetal bovine serum (Cytiva, SH30396.03), non-essential amino acids (Cytiva, SH30238.01), GlutaMAX (Thermo Fisher Scientific, 35-050-061) and penicillin-streptomycin (Corning, MT30002CI). For the cell line expressing EGFP on the outer mitochondrial membrane, the TOM20-EGFP gene was introduced into U2OS cells via lentiviral transduction as described previously^47^. For the cell line expressing EGFP in the mitochondrial matrix, the COX8A-EGFP gene was introduced into U2OS cells using the same procedure. These stable cell lines were maintained under the same conditions and culture media with 5 µg/mL blasticidin added as the selection antibiotics. Prior to imaging, these modified U2OS cells were seeded at a density of 8 × 10^5^ cells on a 35 mm glass-bottom dish with 20 mm micro-well (Cellvis, D35-20-1.5-N) in culture media with 10 mM HEPES added and incubated at 37 °C overnight. For imaging, the dishes were fully filled with culture media and sealed shut using parafilm on the lid.

To image, these sealed dishes were flipped upside down to place the U2OS cells directly under the objective.

#### Live jellyfish preparation

mCherry expressing, transgenic *Clytia hemisphaerica* jellyfish were generated as previously described^48^. Briefly, jellyfish oocytes and sperm were collected, and were micro-injected with a construct driving mCherry under the control of a promoter that expresses the RFamide neuropeptide. To image, jellyfish were embedded in low-melting agarose at a 0.75% concentration in seawater. Animals were then placed in a glass-bottom dish (Willco Wells, Cat. No. GWSB-5040). Once embedded and placed in the dish, the agarose was allowed to solidify for 20 minutes at room temperature. The dish was then flooded with seawater and sealed shut using parafilm on the lid. To image, these sealed dishes were flipped upside down to place the jellyfish directly under the objective. For volumetric imaging, jellyfish were mounted on a glass slide and compressed using a coverslip with clay “feet” on the corners. The edges were then sealed with Vaseline to prevent evaporation. This mounting process fixed the jellyfish in place and minimized movement.

### Live sample imaging

All live sample imaging tasks were performed under ambient conditions. No PH, CO_2_, or temperature control was performed during experiments.

## ACKNOWLEDGEMENTS

We thank Zahid Yaqoob for the loan of the 1.42NA/60× objective lens, Yifei Lu for assistance with the Airyscan experiments, Francisco Sanchez and Ruifeng Hu for their assistance with sample preservation, Tianyu Wang for a helpful discussion on data acquisition system design, and Tianyu Wang and Lei Tian for initial discussions on theory. J.K. was supported by the Picower Postdoctoral Fellowship. This work was partially supported by the National Science Foundation (cooperative agreement EEC-1647837 and award #2449561), the National Institutes of Health (R01NS116139, R01EB029171, R01GM160992, R35GM128859), the Picower Institute for Learning and Memory, and the Freedom Together Foundation.

## AUTHOR INFORMATION

Y.L. current address: Department of Electrical Engineering and Computer Sciences, University of California, Berkeley, CA 94720, USA.

J.Z. current address: School of Biomedical Engineering, University of Oklahoma, Norman, OK 73019, USA.

## Contributions

J.M. and C.L. conceived the projects. C.L. designed and built the system. C.L. wrote and developed the reconstruction algorithm with discussion from B.Z. and Y.L.. Q.L., J.K., and J.Z. prepared experimental samples. C.L. performed imaging experiments with assistance from Q.L. and J.K.. C.L. processed the data. C.L., Q.L., J.K., B.Z., and Y.L. analyzed data. C.L. and J.M. performed simulations. C.L. and J.M. wrote the paper. All authors reviewed the manuscript. J.M., J.N., B.W., and T.B. supervised the project.

## COMPETING INTERESTS

J.M. and C.L. have filed a technology disclosure with Boston University related to the CRM method described in this manuscript.

## ADDITIONAL INFORMATION

Supplementary Information is available for this paper.

## EXTENDED DATA

**Extended Data Fig. 1.**
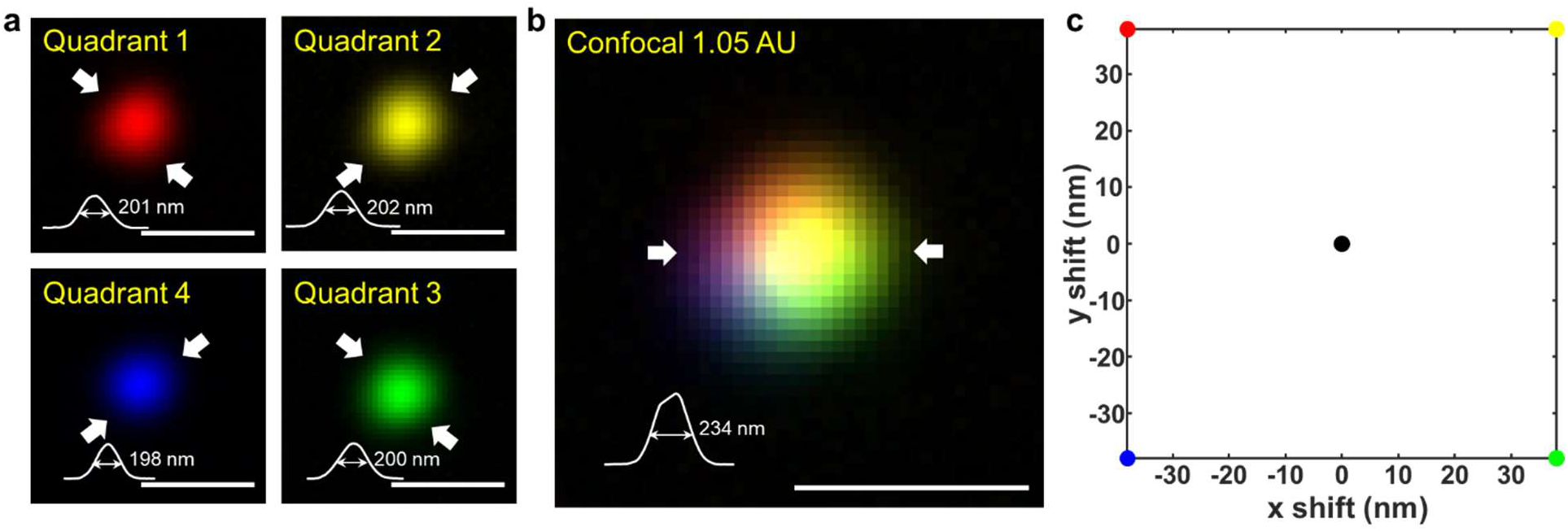
PSF response of CRM system. **a.** PSFs measured from 100 nm fluorescent beads for each detector quadrant. Scale bar: 400 nm. **b.** Confocal PSF (pinhole, 1.05 AU) obtained by summing four quadrants. Each color represents a different quadrant. Scale bar: 400 nm. **c.** Position map of the PSF from each quadrant relative to the confocal PSF. All distance values are converted to the sample plane. The PSF measurement was obtained with a 60×/1.42NA objective. Three independent experiments were performed with similar results.

**Extended Data Fig. 2.**
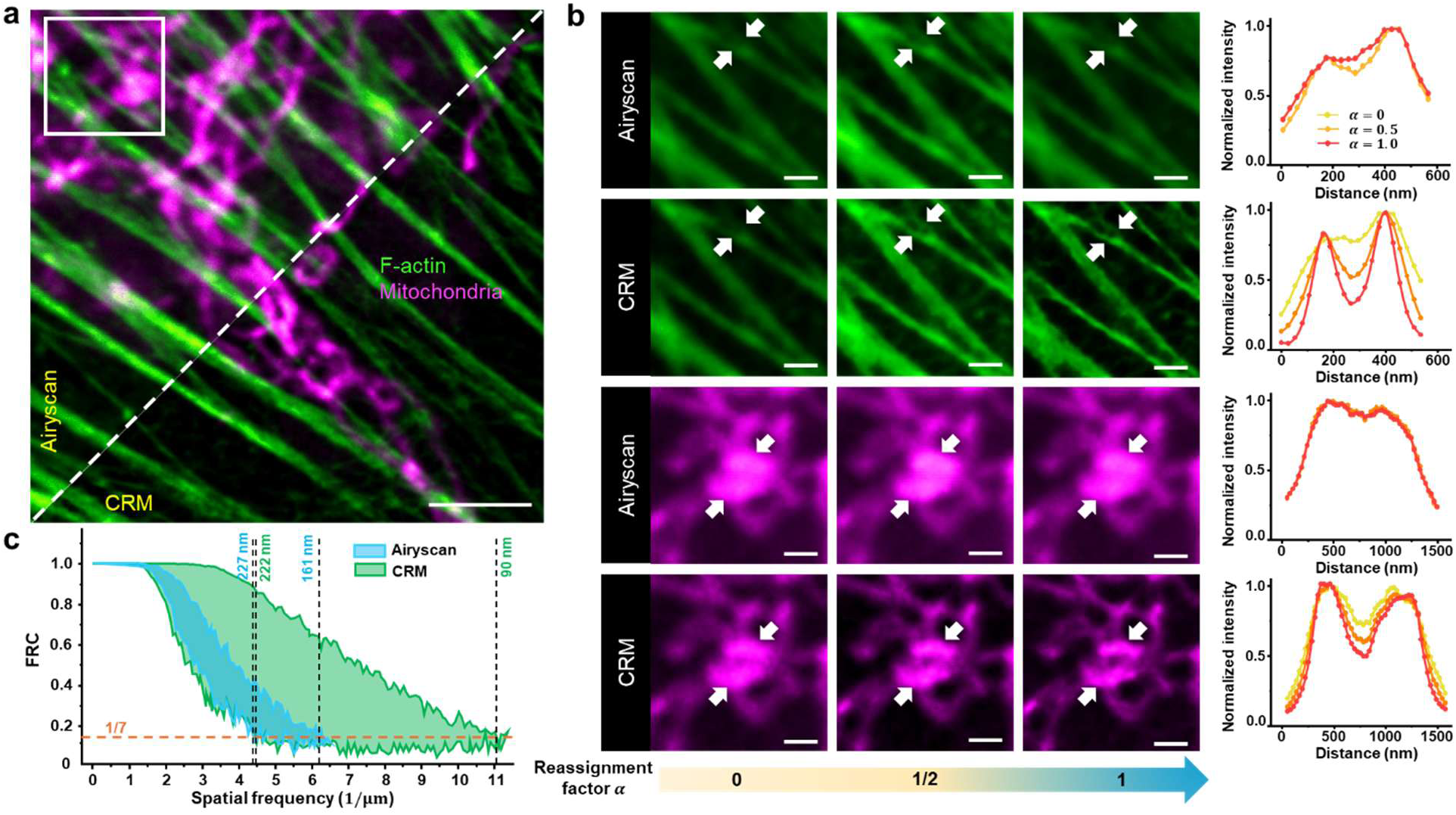
Comparison of effect of reassignment factor on two-color CRM and ISM (Zeiss LSM 980 Airyscan2) resolution**.a.** Two-color images (obtained sequentially) of Alexa Fluor 488 phalloidin-labeled F-actin and MitoTracker Red CMXRos-labeled mitochondria in fixed BPAE cell imaged by Airyscan (left, detection area 1.25 AU) and CRM (right, pinhole 1.05 AU). Scale bar: 3 µm. **b.** Zoomed region (white box) obtained with different reassignment factors (α = 0, 0.5, 1). Intensity profiles across spans indicated by the white arrows are color-coded according to α. Note: Image sharpness increases with α for CRM, whereas it is maximum when α ≈ 0.5 for ISM. Scale bar: 800 nm. **c.** Spatial resolution inferred from FRC analysis in F-actin images, as a function of reassignment factors ranging from 0 to 1. Orange dashed line indicates the conventional FRC threshold used to define resolution. Experiments were repeated three times independently with similar results.

**Extended Data Fig. 3.**
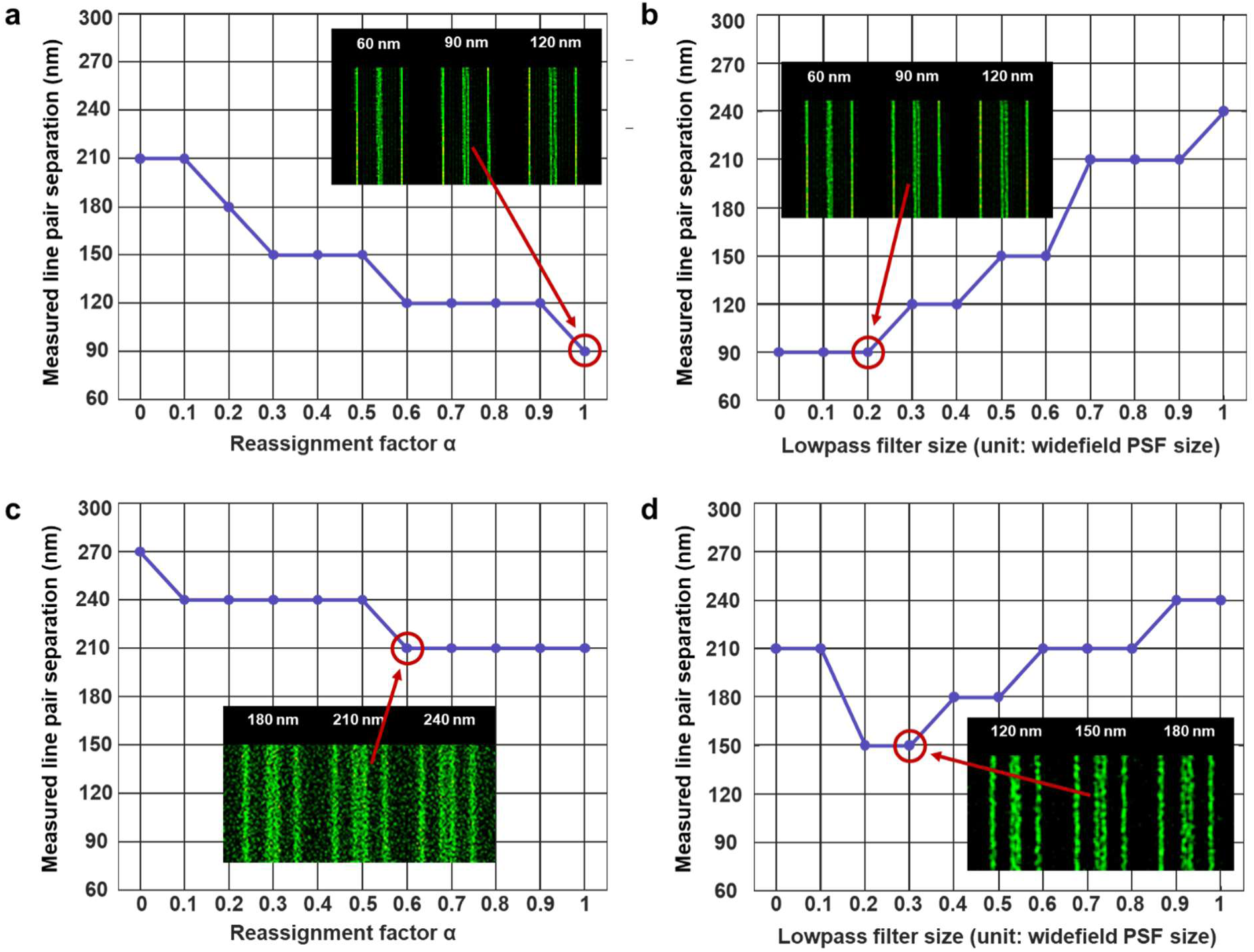
Comparison of reassignment-factor tuning versus lowpass-filter tuning for increased robustness to noise. **a. c.** Reconstruction results as a function of reassignment factor α (no lowpass filter applied to centroid map). **b. d.** Reconstruction results as a function of lowpass filter size (fixed α = 1). Top row, high-SNR data. Bottom row, low-SNR data acquired with 10× shorter pixel dwell time. Insets show representative CRM images of an Argo-SIM resolution target consisting of gradually spaced lines. Under high-SNR conditions, both methods achieve a best resolution of 90 nm, while under low-SNR conditions, lowpass-filter tuning yields a better optimum (150 nm) than reassignment-factor tuning (210 nm), indicating improved robustness to noise. Imaging was performed with a 60×/1.42NA objective. Experiments were repeated three times independently with similar results.

**Extended Data Fig. 4.**
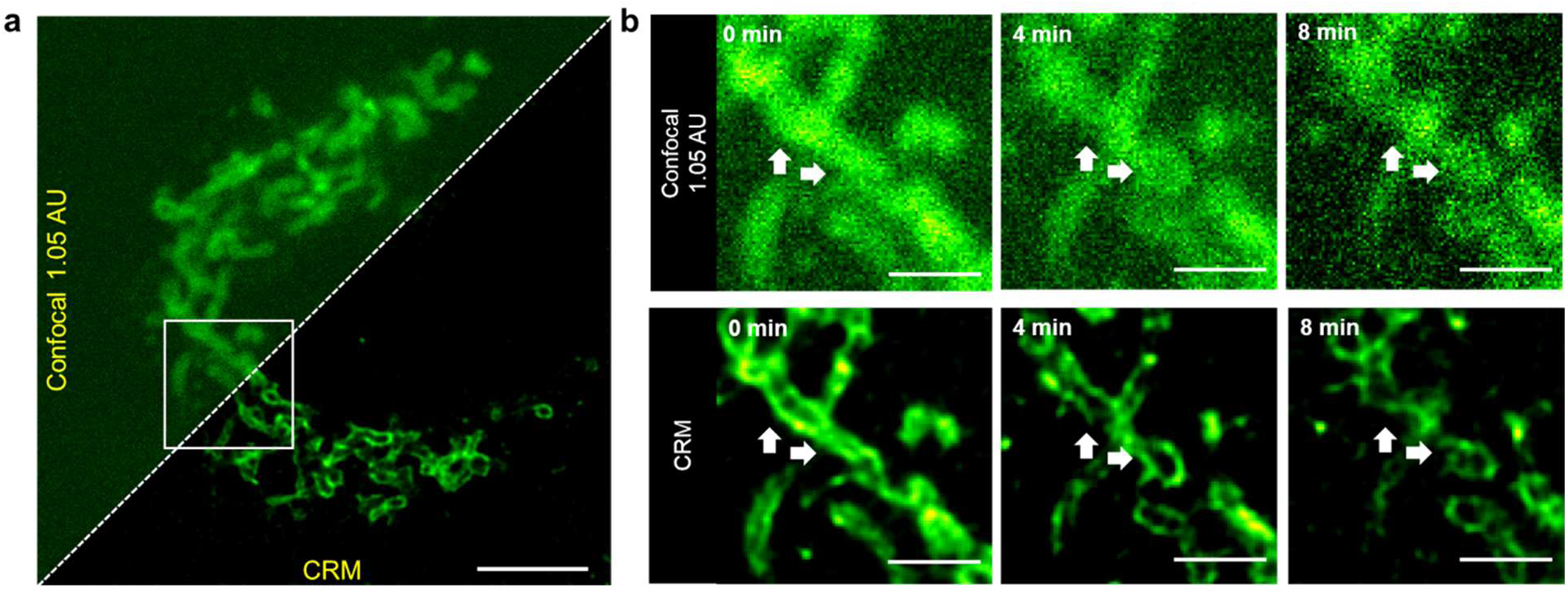
Mitochondrial fission dynamics imaged with CRM. **a.** TOM20-EGFP labelled mitochondria of the same live U2OS cell as in Fig. 4a. Scale bar: 3 µm. **b**. Time-lapse of mitochondrial fission dynamics from the zoomed region (white box) in **a**. White arrows indicate a fission process of a mitochondrion into two. Scale bar: 1 µm. Full video is shown in Supplementary Video 1.

**Extended Data Fig. 5.**
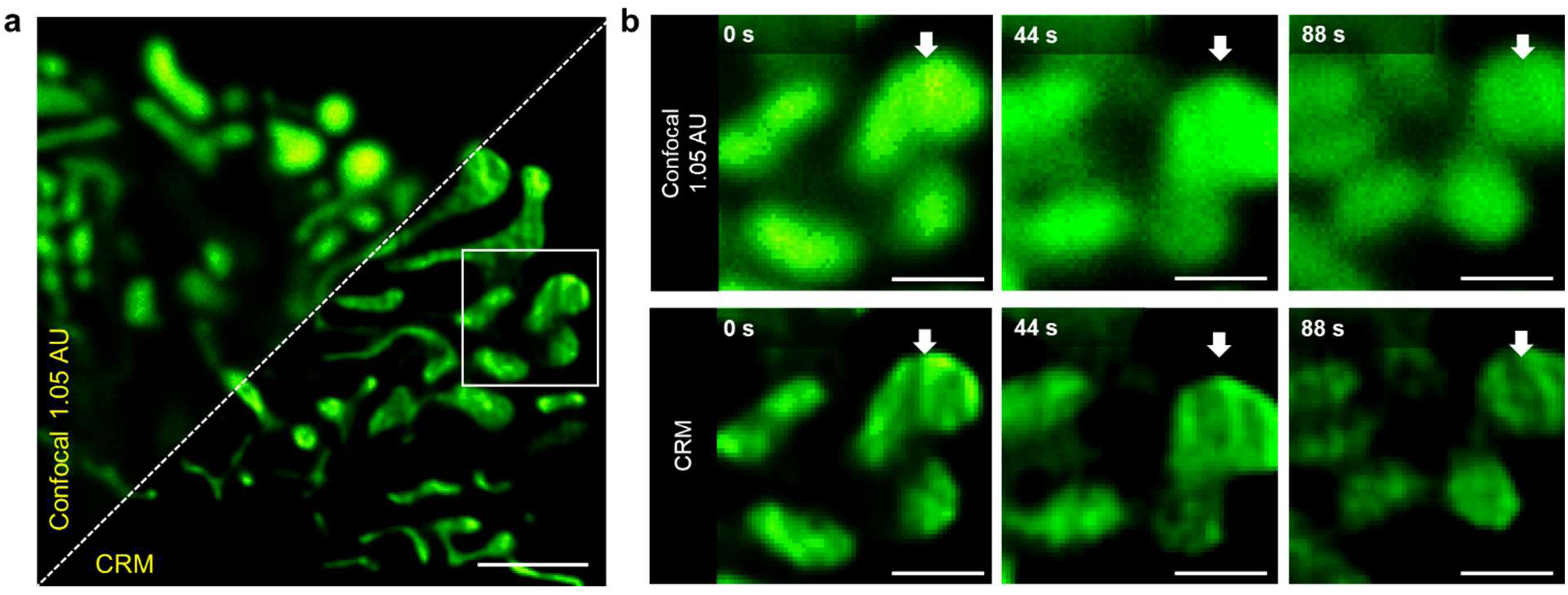
Video-rate (30 Hz) imaging of mitochondrial matrix. **a.** TOM20-EGFP labelled mitochondrial matrix in the live U2OS cell imaged by confocal (summed quadrants, pinhole 1.05 AU) and CRM. Scale bar: 2 μm. **b.** Time-lapse images of mitochondrial matrix dynamics from the zoomed-in region delimited by the white box in **a**. White arrows indicate the internal shape change of a mitochondrion during its interaction with another. Scale bar: 800 nm. The imaging was performed with the 60×/1.42NA objective. The full time-lapse sequence is shown in Supplementary Video 3. Similar imaging experiments were performed three times with similar results.

**Extended Data Fig. 6.**
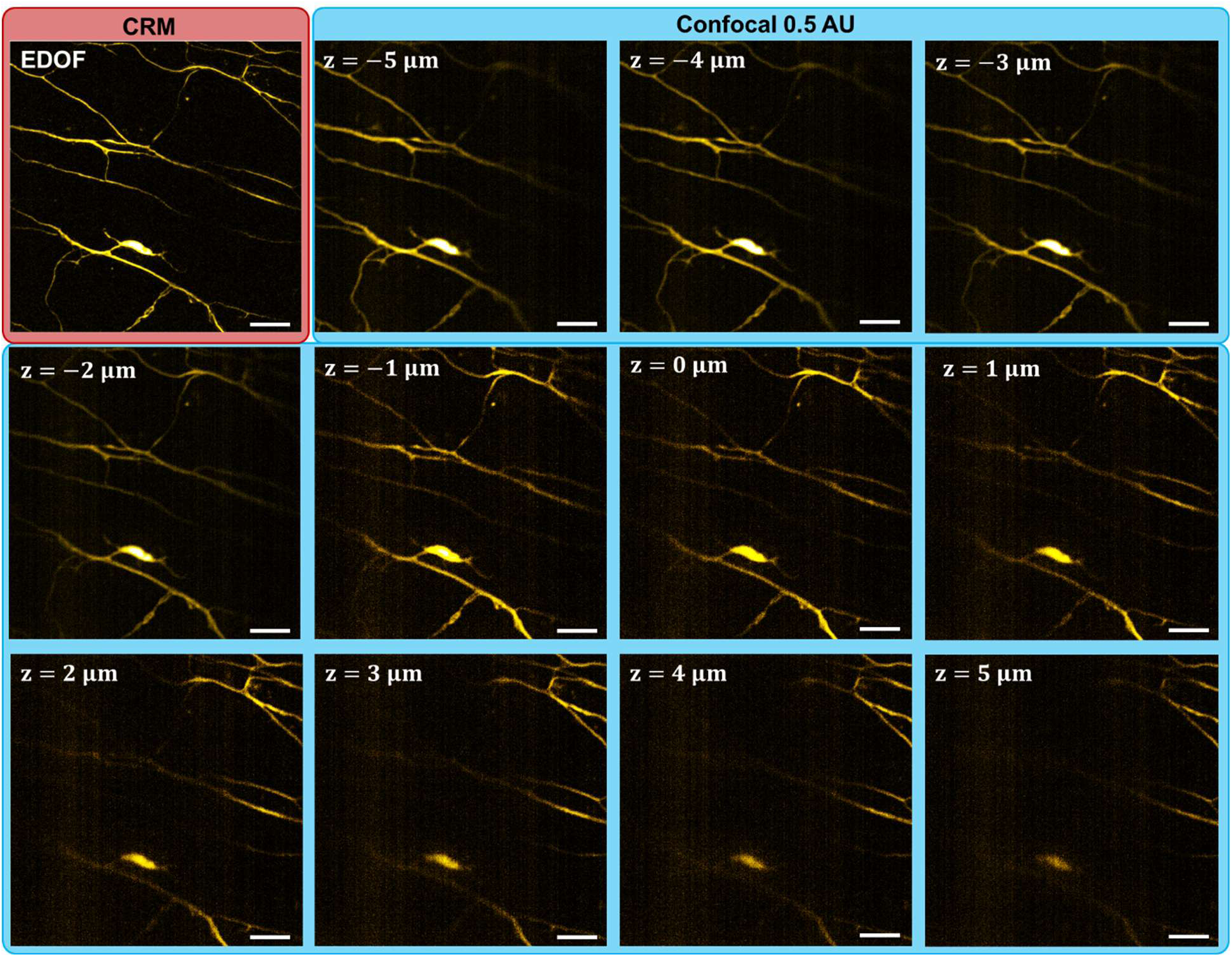
EDOF imaging of jellyfish associated with Fig. 5. Comparison of an EDOF image captured from CRM (pinhole 1.05 AU) and a z-stack captured from conventional confocal microscopy (pinhole 0.5 AU). The smaller confocal pinhole provides improved optical sectioning and serves as the reference for validating the CRM EDOF reconstruction. Scale bar: 3 μm.

## Notes

### Summary of Updates

All figures updated. Methods added. Acknowledgement updated. Author affiliations updated.

